# Benchmarking Peptide Structure Prediction with AlphaFold2

**DOI:** 10.1101/2022.02.17.480937

**Authors:** Alican Gulsevin, Jens Meiler

## Abstract

AlphaFold2 (AF2) is a computational tool developed for the determination of protein structures with high accuracy. AF2 has been used for the modeling of many soluble and membrane proteins, but its performance in modeling peptide structures has not been systematically investigated so far. We benchmarked the accuracy of AF2 in predicting peptide structures between 16 – 60 amino acids using experimentally-determined peptide structures as reference. Our results show that while AF2 can predict the structures of certain peptide scaffolds with RMSD values below 3 Å, it is less successful in predicting the structures of peptides that have kinks, turns, or have extended flexible regions. Further, AF2 had several shortcomings in predicting rotamer recoveries, disulfide bonds, and the lowest RMSD structures based on pLDDT values. In summary, AF2 can be a powerful tool to determine peptide structures, but additional steps may be necessary to analyze and validate the results.

## 1. Introduction

### Peptides: structure and function

Peptides can be loosely defined as polyamides that consist of 2 – 50 amino acids, though this is an arbitrary definition and many molecules accepted to be peptides rather than proteins are larger than this cutoff [1]. Often, the amount of secondary and tertiary structure formed influences whether a polyamide is classified as peptide (less structure) or protein (more structure), in addition to its length. There is a large number of peptides in nature that play important roles as hormones [2,3], antimicrobials [4], and many synthetic peptides are being investigated as drug candidates [5]. In addition to these roles, many toxins are peptides [6]. While flexible, peptides adopt a variety of (temporary) conformations that interact with binding partners and can be sensitive to environmental factors or interactions with lipids or proteins. Typically, solid-state or solution NMR spectroscopy are used for the determination of small peptide structures [7,8]. NMR spectroscopy allows the determination of peptide structures under different experimental conditions, shedding light on how peptides may behave in different environments including different lipid compositions, temperatures, and pH values. Shortcomings of NMR spectroscopy are mostly related to the ambiguities in assignment of signals, which results in an ensemble of structures rather than a single well-defined structure [9]. Considering these challenges, computational methods pose plausible alternatives for the determination of the most stable potentially bioactive conformations of peptides.

### Computational methods for peptide structure determination

There are multiple methods to predict peptide structures including *de novo* folding, homology modeling, molecular dynamics (MD) simulations, and deep-learning-based methods [10–12]. Rosetta *de novo* is based on *de novo* folding principles that insert pre-calculated fragments into a pose and calculate the score for the generated structure [13]. These calculations can be run in solution and in implicit membrane environments that allows the fine tuning of the peptide environment [14]. Typically, thousands of structures are generated, clustered, and analyzed based on predicted energy (score) to determine the likely native-like conformations within the generated ensembles. This method can also be combined with experimental restraints from (NMR) experiments to improve the quality of the results [15–17]. PEP-FOLD3 is a *de novo* folding based method that can be used to model peptides between 5 and 50 amino acids [18]. APPTEST is a protocol that utilizes a neural network architecture, which can be used to model peptides that are between 5 and 40 amino acids [19]. PEPstrMOD utilizes short molecular dynamics simulations to predict peptide structures [20,21]. Calculations can be run in hydrophobic and hydrophilic implicit membranes. One advantage of this method is that it can be applied to model peptides bearing certain non-canonical amino acids or modified termini. In addition to these methods, homology modeling can be used with or without implementing experimental data when there is a homologous peptide or protein structure available [22]. While this method is typically preferred for larger proteins, it was also applied to modeling peptide structures [23]. However, homology modeling requires the presence of template structures with high sequence homology to the target protein. Overall, a computational method that bridges the gaps of the existing computational methods would be useful for the modeling of peptide structures.

### Structure determination with AlphaFold2

The availability of the AF2 source code was a big step for towards the determination of protein structures with high accuracy. AF2 uses deep neural networks trained on known experimental structures to determine protein structures from sequence based on multiple sequence alignments it generates using known protein sequences in combination with the structures of homologous proteins [24]. While AF2 can be used for the modeling of shorter peptides in theory, the benchmark set used to train AF2 excluded shortest peptide structures since the method of determination for these peptides is NMR spectroscopy in general. Some of the relatively poor predictions made by AF2 in CASP14 included protein structures determined by NMR [25], raising the question whether a similar pattern would be observed for flexible peptide structures as well. Therefore, comprehensive benchmarks are necessary to determine the utility of AF2 in modeling peptide structures. Although there are ongoing works on assessing the performance of AF2 in predicting peptide – protein complex structures [26–28], no work focusing solely on small peptide structure determination was published so far. In this work, our aim is to lay the foundation for the use of AF2 to predict the structure of peptides that are between 16 and 60 amino acids, which will prove useful for large scale modeling of peptide structures that are hard to obtain through experimental methods. To achieve this goal, we selected 155 peptides from the Protein Data Bank (PDB) [29] and ran AF2 calculations with these peptides to predict their structures. Next, we compared the predicted and experimental structures to calculate the performance of AF2 in predicting these structures as measured by root mean square deviation (RMSD) and the percentage of recovered rotamers. Finally, we made a case-by-case analysis of the structures that could not be accurately modeled by AF2 to better understand the limitations of the algorithm in modeling peptides.

## 2. Results

### Structure selection and analysis

The structure selection and analysis are described in the Methods section. Briefly, a total of 155 peptides (Supporting Table 1) with experimentally determined structures that included well-defined secondary structure elements were selected. The peptides were split into the following benchmark sets: helical membrane-associated peptides, helical soluble peptides, mixed secondary structure membrane-associated peptides, mixed secondary structure soluble peptides, β-hairpin peptides, and disulfide-rich peptides (DSRP). For each peptide, the lowest-energy NMR structure was compared with all five structures generated by AF2, and the lowest-RMSD pose was used for the analyses. Considering the ambiguities associated with NMR structure determination, an RMSD value less than 3 Å was considered as “good accuracy”, and an RMSD value between 3 and 6 Å was considered as “moderate accuracy” for each structure. An RMSD larger than 6 Å was considered as “poor accuracy”, and the peptides belonging to this group were considered as outliers for each set and investigated further to explain the reason behind their behavior.

**Table 1:**
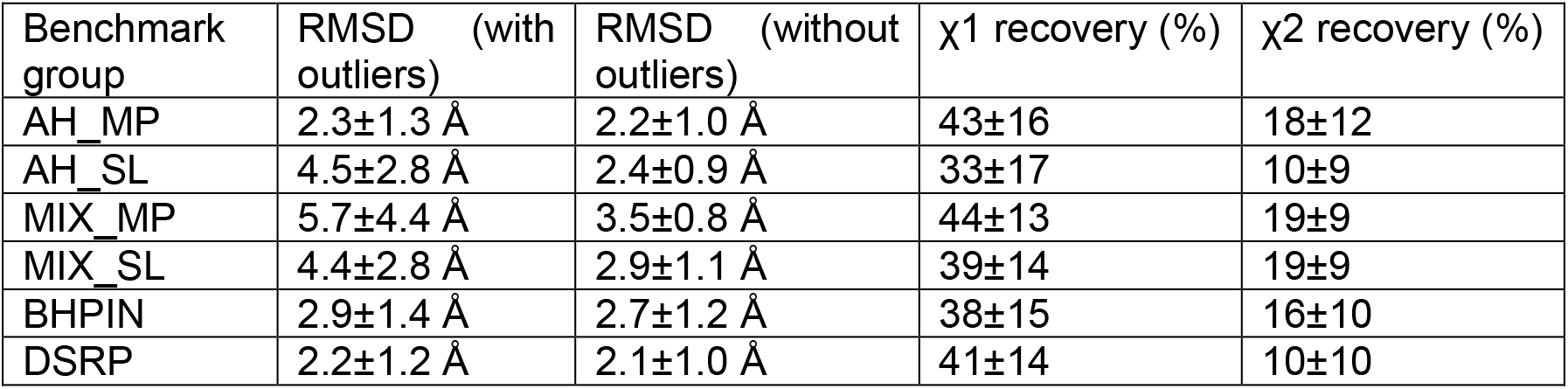
The RMSD values calculated for each benchmark set with and without the outliers included, and the X^1^ and X^2^ recovery percentages calculated for all the structures within the set. All the RMSD values were calculated based on the lowest RMSD value given by the comparison of the five structures generated by AF2 and a reference structure from the PDB. AH_MP stands for helical membrane-associated peptides, AH_SL stands for helical soluble peptides, MIX_MP stands for mixed secondary structure membrane-associated peptides, MIX_SL stands for mixed secondary structure soluble peptides, BHPIN stands for β-hairpin peptides, and DSRP stands for disulfide-rich peptides.

### Helical membrane-associated peptides were predicted with good accuracy and very few outliers

These peptides are defined as peptides that fold into a predominantly helical structure in the presence of a membrane environment. This group covers peptides including transmembrane helices, amphipathic helices, structures with a helix-turn-helix motif, and monotopic helices that do not completely span the membrane. This group was the largest of all the investigated groups, consisting of 67 peptides. The average RMSD calculated for this group of peptides was 2.3±1.3 Å (Table 1). Fifty peptides had RMSD values smaller than 3 Å, fifteen peptides had RMSD values between 3 and 6 Å, and two peptides had RMSD values larger than 6Å (PDB ID: 1SUT and 1VPC, Figure 1, top row). 1SUT was predicted to be unstructured by AF2 whereas it formed a well-defined α-helix in NMR experiments. For 1VPC, the opposite was observed and the residues 78-96 that were unstructured in NMR experiments were predicted to have an α-helical structure by AF2. The helices that had RMSD values within the 3–6 Å range suffered from three things mainly. Some of these helices (1CFG, 1FW5, 1KDL, 1KZ5, 1PEH) had extended helical regions in AF2 calculations, which were coiled in experimental structures. Another group (1FJK, 1IYT, 1LBJ, 1Q2F, 1RG3, 1XC0) had the angle between the two helical regions predicted wrong, which increased the overall RMSD despite accurate prediction of the overall secondary structures. The last group (1EQX, 1KMR, 1MF6) had less well-defined secondary structures compared to the reference structure. As an exception, 1SOL was predicted to have no secondary structure despite the α-helical nature of the reference structure, but the correct angle of the coiled regions resulted in an RMSD lower than 6 Å. When the two outliers were excluded, the calculated RMSD for the helical membrane peptides slightly dropped to 2.2±1.0 Å. The total rotamer recovery rates were 43±16 % for X^1^ and 18±12 % for X^2^ (Table 1).

**Figure 1:**
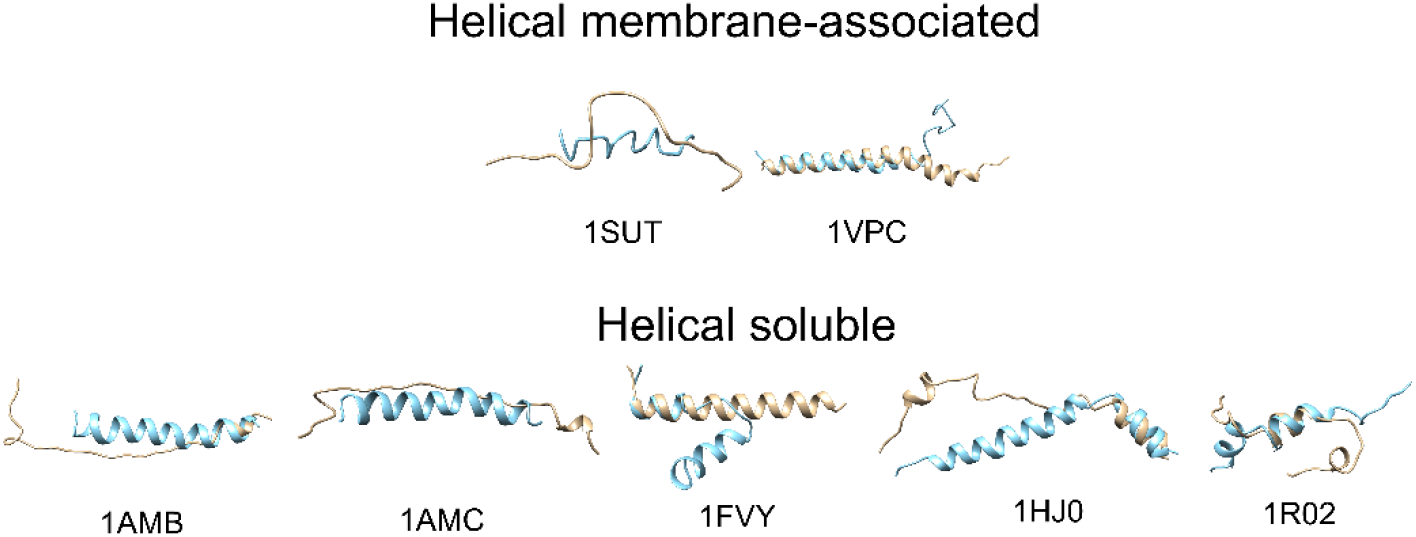
The outlier structures identified for the helical membrane-associated (top row) and helical soluble protein (bottom row) benchmark sets. The beige color stands for the AF2 predicted structures and the blue color stands for the experimental structures.

### Helical soluble peptides had a large number of outliers and performed worse compared to their membrane-associated counterparts

The helical soluble peptide group was defined as α-helical peptides whose structures were not identified in a membrane environment, had no remarks regarding membrane interactions in the original publication, but fulfilled the remaining conditions previously described for helical membrane peptides. Thirteen peptides belonged to this group. The average RMSD calculated for the helical soluble peptides was 4.5±2.8 Å (Table 1). The large average RMSD values and the deviations were caused by the large number of outliers. Six peptides had RMSD values smaller than 3 Å, two peptides had RMSD values between 3 Å and 6 Å, and five peptides had RMSD values larger than 6 Å (PDB ID: 1AMB, 1AMC, 1FVY, 1HJ0, 1R02; Figure 1, bottom row). The reasons for the deviations were similar to that of the helical membrane peptides. The α-helical 1AMB and 1AMC were predicted to be unstructured by AF2. The α-helical regions of 1HJO and 1R02 were also predicted to be partly unstructured (1-30 and 24-33 respectively). 1FVY with its helix-turn-helix motif with a nine amino acid turn region was predicted to be a single α-helix. Without the outliers, the remaining peptides had an average RMSD value of 2.4±0.9 Å. The total rotamer recovery rates were 33±17 % for X^1^ and 10±9 % for X^2^ (Table 1).

### Mixed secondary structure membrane-associated peptides showed the largest variation and RMSD values among all the benchmark sets

The mixed secondary structure peptides were identified to interact with membranes like the helical membrane peptides, but they consisted of more than one secondary structure regions (e.g., multiple α-helices separated by a turn, α/β or α/coil mixed secondary structure, etc.). Thirteen peptides belonged to this group with an average RMSD of 5.7±4.4 Å(Table 1). Three peptides had an RMSD value smaller than 3 Å, six peptides had RMSD values between 3 Å and 6 Å, and four peptides had RMSD values larger than 6 Å (PDB ID: 1CEU, 1GW4, 1PYV, 2JOU; Figure 2, top row). Of the four outliers, 1CEU, 1GW4, and 2JOU had helix-turn-helix angles different than the experimental structures. 1PYV was predicted to be partially unstructured (23-29) that affected the predicted angle between the α-helical regions as well. Even with the outliers removed, the average RMSD only dropped to 3.5±0.8 Å. The total rotamer recovery rates were 44±13 % for X^1^ and 19±9 % for X^2^ (Table 1).

**Figure 2:**
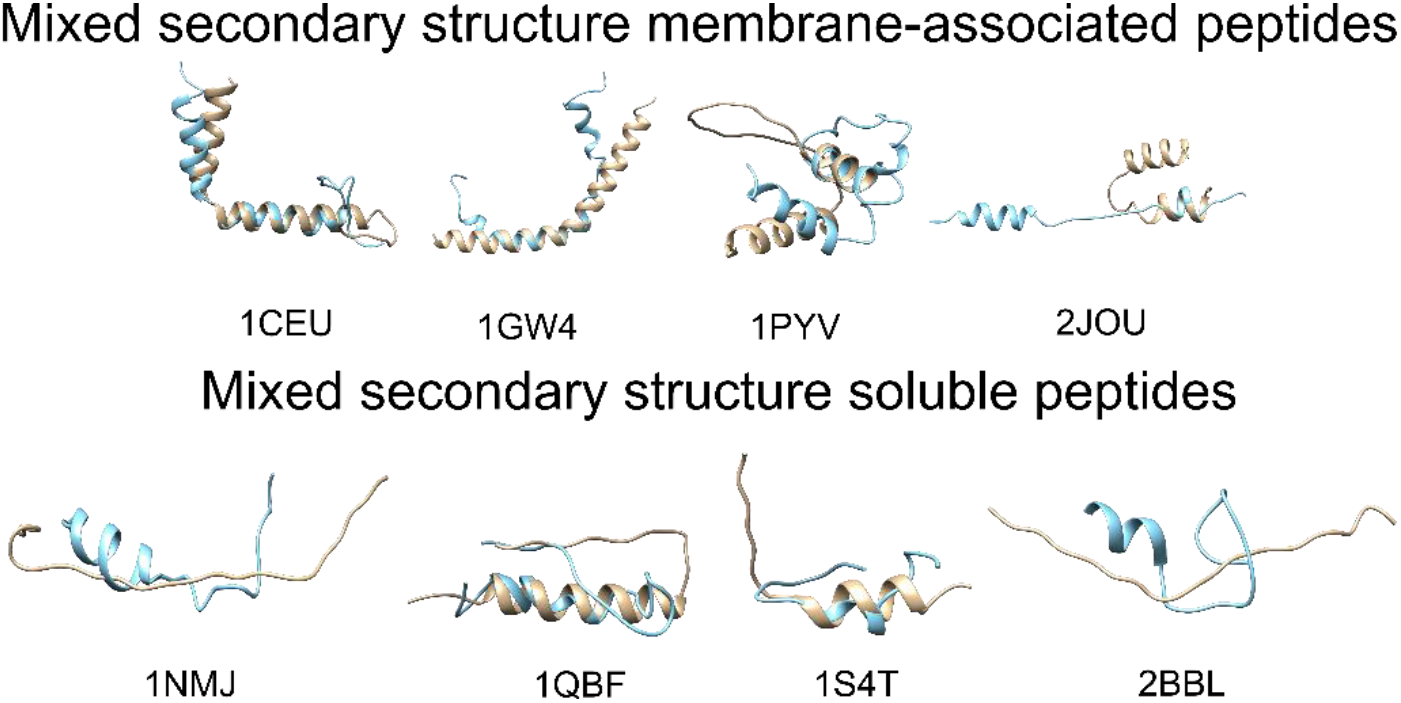
The outlier structures identified for the mixed secondary structure membrane-associated (top row) and mixed secondary structure soluble protein (bottom row) benchmark sets. The beige color stands for the AF2 predicted structures and the blue color stands for the experimental structures.

### Mixed secondary structure soluble peptides had moderate accuracy, which improved upon the removal of outlier peptides

Mixed secondary structure soluble peptides group was defined as peptides that have the same secondary structure properties as their membrane counterparts, but for the peptides whose structures were not identified in a membrane environment. Fourteen peptides belonged to this group with an average RMSD of 4.4±2.8 Å (Table 1). Six peptides had an RMSD value smaller than 3 Å, four peptides had RMSD values between 3 and 6 Å, and four peptides had RMSD values larger than 6 Å (PDB ID: 1NMJ, 1QBF, 1S4T, 2BBL; Figure 2, bottom row). 1NMJ and 2BBL were predicted to be completely unstructured in AF2 calculations. 1QBF had the 13-22 region α-helical in AF2 calculations, but this region was predicted to be a coil region in the NMR structure. 1S4T had a coiled N-terminus consisting of ten amino acids, but the angle of this region was predicted wrong by AF2. The average RMSD values calculated without the outliers was 2.9±1.1 Å. The total rotamer recovery rates were 39±14 % for X^1^ and 19±9 % for X^2^ (Table 1).

### β-hairpin peptides were predicted with good accuracy for both stapled and non-stapled peptides

The β-hairpin peptides group includes peptides that have a single β-hairpin motif. Members of this group may or may not be stapled by the presence of a disulfide bond. There was a total of 22 peptides in this group (18 stapled and 4 non-stapled) with an average RMSD of 2.9±1.4 Å (Table 1). Thirteen peptides had an RMSD value smaller than 3 Å, eight peptides had RMSD values between 3 Å and 6 Å, and one peptide had an RMSD value larger than 6Å (PDB ID: 1G04; Figure 3, left panel). For the peptides with moderate accuracy, the β-hairpin motif was predicted accurately, but the curvature of the tip of the hairpin varied from the experimental measurements in some cases. 1G04 was predicted by AF2 to have an α-helical structure with a 10-12 amino acid long coiled C-termini. The average RMSD without the outlier was 2.7±1.2 Å. The total rotamer recovery rates were 38±15 % for X^1^ and 16±10 % for X^2^ (Table 1).

**Figure 3:**
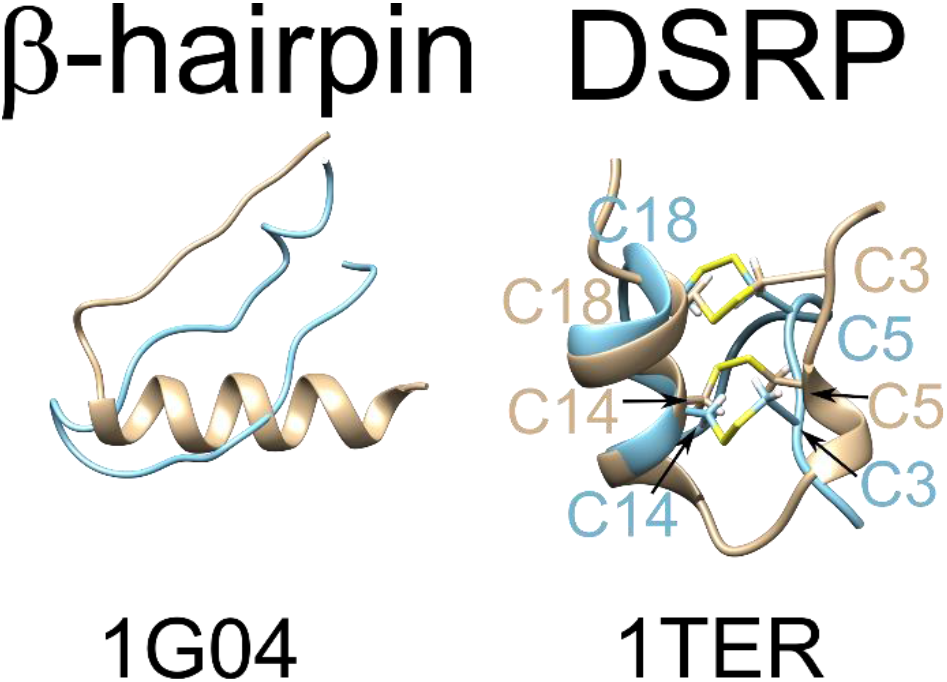
The outlier structures identified for the β-hairpin (left) and DSRP (right) benchmark sets. The beige color stands for the AF2 predicted structures and the blue color stands for the experimental structures. The cysteines contributing to the disulfide bonds of the experimental and predicted structures are labeled explicitly in the color of the corresponding peptide structure.

### Disulfide-rich peptide structures were predicted with high accuracy, but the disulfide bonding patterns deviated from the reference structures

Disulfide-rich peptides (DSRP) were defined in the context of this work as any peptide that had two or more disulfide bonds. This group included toxin peptides such as α-conotoxins, β-hairpins cyclized by multiple disulfide bonds, and some hormone peptides. A total of 26 DSRPs were used for the analyses, and the average RMSD value of 2.2±1.2 Å was the lowest among all groups (Table 1). 20 peptides had an RMSD value smaller than 3 Å, five peptides had RMSD values between 3 and 6 Å, and one peptide had RMSD values larger than 6Å (PDB ID: 1TER)(Figure 3, right panel). The deviation of the outlier was caused by wrong disulfide pairing (3,18 – 5,14 instead of 5,18 – 3,14). The average RMSD value without the outlier was 2.1±1.0 Å. The total rotamer recovery rates were 41±14 % for X^1^ and 10±10 % for X^2^ (Table 1).

Next, we checked the disulfide connectivity patterns to investigate whether the high accuracy AF2 showed for the DSRP benchmark set is due to accurate assignment of disulfide bonds. Surprisingly, 35% of the tested peptides had one or more cysteines either without a disulfide bonding pair or forming disulfide bonds with a cysteine different than observed in the reference structure (Supporting Table 2). A closer inspection of the five structures generated for each sequence showed that AF2 may produce results with consistent and accurate representation of the disulfide bonds in some cases (Figure 4A), whereas in other cases it makes predictions that involve different disulfide bonding patterns (Figure 4B) or predictions that lack disulfide bonds in some cases (Figure 4C).

**Figure 4:**
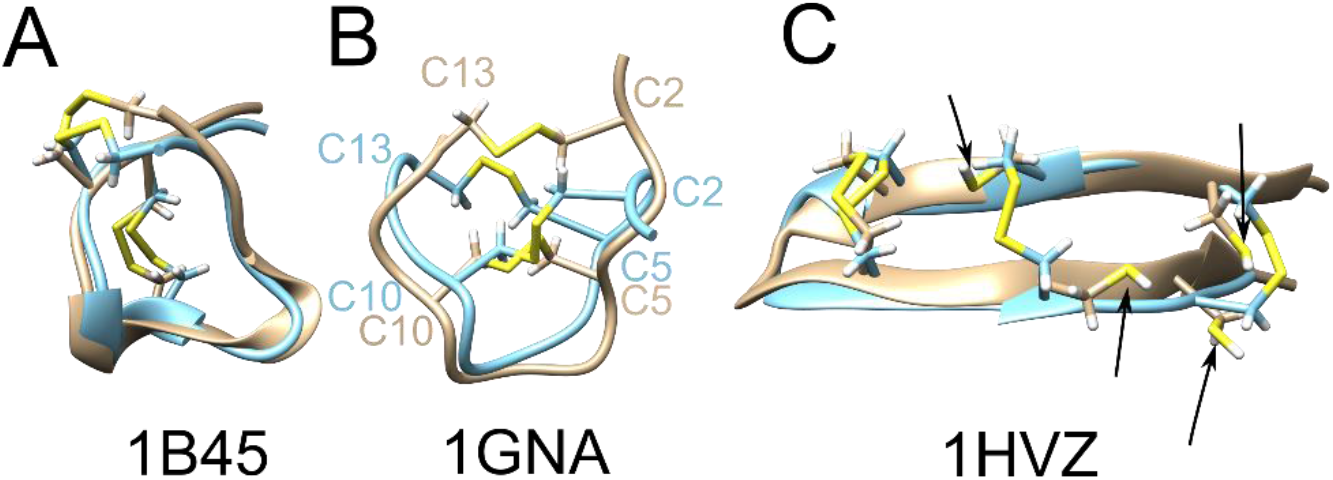
Alignment of select predicted (beige) and experimental (blue) DSRPs. The disulfide bonds are shown explicitly. (A) Example to disulfide bonds consistent with the experimental structure, (B) example to an alternative predicted disulfide bonding pattern, (C) examples to lack of disulfide bond formation between pairs that form disulfide bonds in experimental structures. Free cysteines are shown with black arrows.

**Table 2:** The RMSD values calculated for each benchmark set with and without the outliers included, and the X^1^ and X^2^ recovery fractions calculated for all the structures within the set when only the top-ranking structure was used for the RMSD calculations. AH_MP stands for helical membrane-associated peptides, AH_SL stands for helical soluble peptides, MIX_MP stands for mixed secondary structure membrane-associated peptides, MIX_SL stands for mixed secondary structure soluble peptides, BHPIN stands for β-hairpin peptides, and DSRP stands for disulfide-rich peptides.

| Benchmark group | RMSD (with outliers) | RMSD (without outliers) | $\chi_1$ recovery (%) | $\chi_2$ recovery (%) |
| --- | --- | --- | --- | --- |
| AH_MP | 2.8±1.9 Å | 2.3±1.0 Å | 44±18 | 19±14 |
| AH_SL | 4.9±3.2 Å | 2.5±0.9 Å | 35±19 | 12±10 |
| MIX_MP | 6.8±4.4 Å | 4.0±1.0 Å | 44±18 | 21±14 |
| MIX_SL | 5.4±3.7 Å | 3.4±1.3 Å | 36±11 | 17±10 |
| BHPIN | 3.5±2.2 Å | 3.0±1.2 Å | 39±13 | 14±09 |
| DSRP | 2.4±1.2 Å | 2.3±1.0 Å | 40±14 | 12±10 |

### There is no correlation between the predicted rank and prediction accuracy

A standard AF2 run generates five structures as output. These structures are ranked by the program based on their predicted local distance difference test (pLDDT) values, which is a metric that estimates how well a predicted structure would agree with an experimental structure based on backbone carbon coordinates [24]. Based on these values, the lowest-ranking structure is considered to be the representative structure of the given sequence. However, this assumption may not always be true. All five AF2 structures generated were compared with the corresponding reference structure in this study to allow better structural sampling. In theory, the structures with lower ranks should give the lowest RMSD values in our calculations because they are considered to be have higher confidence by AF2. To test whether this assumption is true, we calculated the percentages of each rank that gave the lowest RMSD for the particular structure. Based on these calculations, there was no correlation between the rank assigned by AF2 and the structure that gave the lowest RMSD. Only 13% of the lowest-RMSD structures came from structures that had a rank of 0 (best rank). For the remaining predictions, 26%, 18%, 24%, and 19% came from ranks 1, 2, 3, and 4 respectively. Therefore, our results suggest that the pLDDT metric that can be used to assess globular protein structures is not a meaningful metric to classify peptide conformations generated by AF2. Comparison of the cases where we calculated the lowest RMSD including all five structures generated by AF2 (Table 1) and only based on the top-ranking structure (Table 2) clearly shows that the latter approach causes an increase in the RMSD values calculated and the number of outliers, though with little effect on X^1^ and X^2^ recovery percentages.

### Limitations of the study

The first limitation of the study is caused by the intrinsic flexibility of many peptides in general. Peptides can have highly flexible regions including coils or turns that may result in multiple conformations for the same structure. NMR structures typically consist of an ensemble of conformations, which makes an exact comparison between the predicted and experimental structures challenging. In order to address this point, predictions with RMSD values lower than 3Å were loosely considered good predictions, and all the tested benchmark sets with the exception of mixed secondary structure soluble peptides had RMSD values below 3Å when the outliers were excluded.

Another point related to flexibility is that alternative low-energy conformations to the structures identified by NMR may exist. Especially with helices that have multiple domains connected by turns or coils, the structures captured by experimental methods may represent only one of the multiple conformations the peptides. From that perspective, the structures predicted by AF2 may not necessarily be wrong, but they may simply correspond to an alternative conformation of the peptide. A study aimed at comparing the quality of AF2-predicted structures with experimental structures determined by NMR in solution identified that structures predicted by AF2 can be more energetically stable and therefore feasible compared to NMR structures in some cases [30]. More systematic studies that focus on peptide dynamics through experimental and computational methods will be necessary to understand whether the AF2-predicted peptide structures that showed great variation from the experimental predictions are likelier to represent the actual structure of these peptides.

Finally, some peptides may fold into different secondary structures depending on the environmental conditions including pH, temperature, and the presence of a membrane. Because the structures from NMR experiments are obtained with a single set of experimental parameters, AF2 predictions may represent alternative conformations of the peptides that could have been obtained under different conditions. This may particularly be issue when AF2 over- or underestimates the secondary structure of peptides, which may indeed be more or less disordered depending on these environmental conditions.

## 3. Discussion

The helical membrane peptide protein structures were predicted overall with good success with the exception of a relatively small number of peptides in each benchmark set. There was no single reason for the deviations caused by the outliers, but the inability of AF2 to predict the secondary structure of some peptides and the inaccurate prediction of the kink angles accounted for most deviations. A similar pattern was observed for the mixed secondary structure membrane and soluble peptides whereby large RMSD values were caused by inaccurate prediction of the coiled regions that connect multiple domains of these peptides. The average RMSD values of 2.2 Å and 2.4 Å calculated for the helical membrane-associated and soluble peptides without the outliers were within the good accuracy range of the calculations, possibly improved because the multitude of the members of these sets were single helical structures, which were predicted with better accuracy than the cases whereby a turn or kink region was present.

The mixed secondary structure peptides were also predicted with varying accuracy. In the case of the membrane-associated peptides, deviations were caused mostly by the errors in prediction of the angles between α-helical regions, but the secondary structure predictions were overall accurate with few exceptions. For the soluble peptides, however, three out of four outliers were predicted to have no secondary structure partly or completely by AF2. On the other hand, the RMSD values show that removal of the outliers yielded RMSD values of 3.5 Å for the membrane-associated peptides and 2.9 Å for the soluble peptides, indicating that even when the secondary structure is predicted correctly, the exact geometry between the different structural domains may not be accurate.

For β-hairpin peptides that had an RMSD value of 2.7 Å without the outlier, the peptides that had the largest RMSD value was predicted to have a different secondary structure, but for the remaining peptides the deviations from the experimental structures were mostly caused by the differences in the geometry of the tip of the hairpin structure. While this may be related to AF2’s inability to capture the right geometry of these flexible regions, it is also possible that the NMR structures were not solved under experimental conditions that would be more consistent with the geometries calculated by AF2. Overall, AF2 was able to accurately predict the geometries of majority of these structures with RMSDs lower than 3.0 Å.

Finally, DSRPs had the highest degree of accuracy among the tested peptides, potentially thanks to their constrained structures due to the presence of multiple disulfide bonds. These structures also had regions with well-defined secondary structures that make predictions easier for AF2 compared to coiled regions where multiple degrees of freedom are present. The RMSD values were the lowest for this group with or without the outlier included (2.2 Å and 2.1 Å respectively). On the other hand, the results demonstrated that the lowest-RMSD AF2 predictions had different or no disulfide bonding patterns for 35% of the peptides even in cases where peptide scaffolds were accurately predicted. This suggests that special care must be shown when predicting DSRP structures that may have alternative disulfide bonding patterns, and the use of additional tools may be necessary to remodel the disulfide bonds following AF2 runs.

Poor rotamer recovery percentages were observed for all the tested peptide sets, whereby the X^1^ values were all lower than 50 % and the X^2^ values were mostly lower than 20 %. This is surprising considering the overall success of AF2 with globular proteins for which the calculated rotamer recoveries were over 80 % in another study [31]. The reason for this discrepancy may be that the investigated peptides have mostly solvent-exposed residues with few buried amino acids due to the small and extended structures of these peptides that prevent close interactions between the different domains. We looked for correlations between the calculated rotamer recovery percentages and parameters like RMSD, solvent accessible surface area, and average per peptide pLDDT for each benchmark set (data not shown), but we found no correlation between these parameters and rotamer recovery percentages. Considering the large deviation from the case of globular proteins, additional tools may be necessary to optimize the side chain conformations of the peptides generated through AF2.

Lastly, our analysis shows that pLDDT, which is the main metric used by AF2 to rank the generated structures is not a good measure of whether a peptide structure is accurately predicted. Only 13% of the lowest-RMSD structures came from the top-ranking structures in our calculations, and there was no correlation among the other estimated ranks and the calculated RMSD values. Further, when RMSD values were calculated only for the top-ranking structures, the RMSD values in all groups increased between 0.2 and 1.1 Å with an increase in the number of outliers in each group. The largest effect was observed for the mixed secondary structure benchmark groups (~1.0 Å with outliers and ~0.5 Å without outliers), suggesting that this shortcoming is a significant problem especially when the system has flexible regions that increase the number of conformations that can be adopted by the peptide but can also cause deviations based on the extent of the secondary structures predicted for the system. A study investigating multiple aspects of protein folding with AF2 involved testing of pLDDT values for proteins of varying lengths, which showed that shorter sequences tend to have larger pLDDT values in general [32]. This may reduce the utility of this metric for small peptide structures by narrowing the gap between good and bad predicted structures. In the absence of a clear selection criterion for the AF2-generated structures, it may be necessary to increase the number of output structures generated and use approaches such as clustering to narrow down the conformations that are sampled more frequently or show more consistent patterns to select the accurate structure.

All these data raise the question of whether the issues regarding the shortcomings of AF2 identified in this study are unique to peptides, or are general issues associated with AF2 making predictions on structures on which it was not trained. Ever since the release of AF2, the focus of the benchmark studies has been what AF2 is capable of doing with little focus on its shortcomings. Our work presents a glimpse into limiting factors that should be considered when modeling peptides, but similar studies that investigate these elements in globular proteins are also necessary in the future.

## 4. Conclusions

AF2 can be used for the modeling of peptide structures smaller than 60 amino acids if the target peptide is anticipated to have a well-defined secondary structure and lacks multiple turn or flexible regions that may assume different conformations. AF2 is particularly successful in the prediction of helical membrane-associated peptides and DSRPs but has reduced accuracy in the cases with extended coiled or flexible regions. Even for the DSRPs that had low RMSDs, issues with disulfide bonding patterns may result in errors in modeling peptides. The poor rotamer recovery rates calculated for the benchmarked peptides suggests that additional tools may be necessary to refine the peptide structures generated by AF2. In addition, AF2 predictions showed no correlation between the predicted ranks and prediction accuracy, therefore alternative metrics to select from the AF2-generated structures may be necessary. Overall, use of AF2 for peptide structure prediction will require development of additional metrics and controls to improve their accuracy.

## 5. Materials and Methods

### Structure selection criteria

A PDB search was done to select peptides smaller than 60 amino acids whose structures were determined through solid-state or solution NMR. Peptides cyclized through their backbones, that have floppy termini, predominantly lacking a secondary structure, lacking side chain confirmations, and peptides bearing non-canonical amino acids were excluded from the set. For the peptides with multiple reported confirmations in different media, the PDB entry that had the most secondary structure was selected if the confirmations were similar, and the entry was excluded if the secondary structure elements showed variability among the different entries. A total of 155 peptides fit these criteria and were used for the benchmark calculations. The benchmark set was divided into further groups to facilitate analysis of the results. These groups are helical membrane peptides, helical soluble peptides, mixed secondary structure membrane peptides, mixed secondary structure soluble peptides, β-hairpin peptides, and disulfide-rich peptides (DSRP).

### AF2 calculations

AF2 calculations were run locally in the monomer mode starting from the sequences of each entry as extracted from the PDB. Terminal modifications were removed from the sequence files prior to the calculations. For each entry, five structures were generated with AF2 and the best-ranking structure according to the LDDT scores was selected as the representative pose for the entry. All the analyses were done with these representative poses.

### RMSD calculations

RMSD values were calculated as the average root mean squared distance between the coordinates of the helix Cα of all five AF2-generated structures and the lowest-energy corresponding reference structure from the PDB. The structures were aligned based on Cα coordinates to eliminate translational differences between the reference and AF2 coordinates. Only the lowest-RMSD AF2 structure was used for the calculation of the average RMSD values. sFor each benchmark set, the RMSDs were calculated twice: one with all the entries in the set, another with the outliers removed to measure how well the good and moderate structures are predicted.

### Rotamer recovery calculations

Rotamer recovery rate calculations were done using an in-house script utilizing BioPython. The peptides PDB structures were used as the reference structures for the comparisons and the lowest-RMSD AF2 structure was used as the decoy structure. The X^1^ and X^2^ angles of each residue of the decoy structures were compared with that of the native structures, and residues with rotamer matches within 25° were considered to be recovered.

## Author Contributions

Conceptualization, A.G.; methodology, A.G.; software, A.G.; validation, A.G.; formal analysis, A.G.; investigation, A.G.; resources, J.M.; data curation, A.G.; writing— original draft preparation, A.G.; writing—review and editing, A.G., J.M.; visualization, A.G.; supervision, J.M.; project administration, J.M.; funding acquisition, J.M. All authors have read and agreed to the published version of the manuscript.

