## Supporting Tables for "Benchmarking Peptide Structure Prediction with AlphaFold2"

### Supporting Information

#### Supporting Tables

Supporting Table 1: List of all the PDB entries used for the benchmark set and the group they were assigned to. AH_MP stands for helical membrane-associated peptides, AH_SL stands for helical soluble peptides, BHPIN stands for β-hairpin peptides, DSRP stands for disulfide-rich peptides, MIX_MP stands for mixed secondary structure membrane-associated peptides, and MIX_SL stands for mixed secondary structure soluble peptides.

| Name | Benchmark group |
|---|---|
| 1A11 | AH_MP |
| 1ALE | AH_MP |
| 1ALF | AH_MP |
| 1B9U | AH_MP |
| 1BDE | AH_MP |
| 1BTQ | AH_MP |
| 1BTR | AH_MP |
| 1BTS | AH_MP |
| 1BTT | AH_MP |
| 1CFG | AH_MP |
| 1CKW | AH_MP |
| 1CKX | AH_MP |
| 1D7N | AH_MP |
| 1DEP | AH_MP |
| 1DSJ | AH_MP |
| 1DTC | AH_MP |
| 1EMZ | AH_MP |
| 1EQX | AH_MP |
| 1FI0 | AH_MP |
| 1FJK | AH_MP |
| 1FW5 | AH_MP |
| 1HLL | AH_MP |
| 1HO2 | AH_MP |
| 1IBO | AH_MP |
| 1IYT | AH_MP |
| 1JAV | AH_MP |
| 1KDL | AH_MP |
| 1KMR | AH_MP |
| 1KZ0 | AH_MP |
| 1KZ2 | AH_MP |
| 1KZ5 | AH_MP |
| 1KZT | AH_MP |
| 1KZV | AH_MP |

| | |
|---|---|
| 1LBJ | AH_MP |
| 1LYP | AH_MP |
| 1MF6 | AH_MP |
| 1MP6 | AH_MP |
| 1ODP | AH_MP |
| 1OEF | AH_MP |
| 1OT0 | AH_MP |
| 1P0G | AH_MP |
| 1P0J | AH_MP |
| 1P0L | AH_MP |
| 1P0O | AH_MP |
| 1P5K | AH_MP |
| 1P5L | AH_MP |
| 1PEH | AH_MP |
| 1PEI | AH_MP |
| 1Q2F | AH_MP |
| 1QFA | AH_MP |
| 1R7C | AH_MP |
| 1RG3 | AH_MP |
| 1RPV | AH_MP |
| 1SOL | AH_MP |
| 1SUT | AH_MP |
| 1T51 | AH_MP |
| 1T52 | AH_MP |
| 1T5Q | AH_MP |
| 1VPC | AH_MP |
| 1XC0 | AH_MP |
| 1XNL | AH_MP |
| 1XOO | AH_MP |
| 2BP4 | AH_MP |
| 2DTB | AH_MP |
| 2GLG | AH_MP |
| 2MAG | AH_MP |
| 2NR1 | AH_MP |
| 1AMB | AH_SL |
| 1AMC | AH_SL |
| 1D1F | AH_SL |
| 1FVY | AH_SL |
| 1HJ0 | AH_SL |
| 1J5B | AH_SL |
| 1JZP | AH_SL |
| 1LB0 | AH_SL |
| 1LCX | AH_SL |

| | |
|---|---|
| 1M23 | AH_SL |
| 1P82 | AH_SL |
| 1R02 | AH_SL |
| 2RMF | AH_SL |
| 1A1P | BHPIN |
| 1B03 | BHPIN |
| 1E0Q | BHPIN |
| 1EGT | BHPIN |
| 1FGD | BHPIN |
| 1FGE | BHPIN |
| 1G04 | BHPIN |
| 1HRL | BHPIN |
| 1IM7 | BHPIN |
| 1KB7 | BHPIN |
| 1KB8 | BHPIN |
| 1LFC | BHPIN |
| 1MPV | BHPIN |
| 1NIL | BHPIN |
| 1NIM | BHPIN |
| 1NJ0 | BHPIN |
| 1O8Y | BHPIN |
| 1O8Z | BHPIN |
| 1PAK | BHPIN |
| 1PAN | BHPIN |
| 1PAO | BHPIN |
| 1QX9 | BHPIN |
| 1B45 | DSRP |
| 1CNL | DSRP |
| 1DG2 | DSRP |
| 1EDP | DSRP |
| 1EI0 | DSRP |
| 1G2G | DSRP |
| 1GNA | DSRP |
| 1GNB | DSRP |
| 1HP9 | DSRP |
| 1HVZ | DSRP |
| 1I8E | DSRP |
| 1IEN | DSRP |
| 1KFP | DSRP |
| 1KWD | DSRP |
| 1KWE | DSRP |
| 1MII | DSRP |
| 1MXN | DSRP |

| | |
|---|---|
| 1PG1 | DSRP |
| 1QMW | DSRP |
| 1RPB | DSRP |
| 1S6W | DSRP |
| 1TER | DSRP |
| 1UL2 | DSRP |
| 1V6R | DSRP |
| 1XGA | DSRP |
| 1XGB | DSRP |
| 1BH1 | MIX_MP |
| 1BL1 | MIX_MP |
| 1CEU | MIX_MP |
| 1FAC | MIX_MP |
| 1GW3 | MIX_MP |
| 1GW4 | MIX_MP |
| 1HOD | MIX_MP |
| 1HZN | MIX_MP |
| 1IBN | MIX_MP |
| 1L4T | MIX_MP |
| 1PYV | MIX_MP |
| 2DWF | MIX_MP |
| 2JOU | MIX_MP |
| 1BBA | MIX_SL |
| 1CQ0 | MIX_SL |
| 1D0W | MIX_SL |
| 1DPK | MIX_SL |
| 1DU1 | MIX_SL |
| 1FDF | MIX_SL |
| 1MEQ | MIX_SL |
| 1NMJ | MIX_SL |
| 1QBF | MIX_SL |
| 1S4T | MIX_SL |
| 1V1D | MIX_SL |
| 2BBL | MIX_SL |
| 2GP8 | MIX_SL |
| 2OOP | MIX_SL |

Supporting Table 2: The list of the DSRP benchmark set members and information on whether the lowest-RMSD structure formed the same disulfide bonds as the reference structure.

| ID | Disulfide Consistent with Reference? |
|---|---|
| 1B45 | Yes |
| 1CNL | Yes |

| | |
|---|---|
| 1DG2 | Yes |
| 1EDP | Yes |
| 1EI0 | Yes |
| 1G2G | Yes |
| 1GNA | No |
| 1GNB | No |
| 1HP9 | Yes |
| 1HVZ | No |
| 1I8E | No |
| 1IEN | Yes |
| 1KFP | Yes |
| 1KWD | Yes |
| 1KWE | Yes |
| 1MII | No |
| 1MXN | No |
| 1PG1 | Yes |
| 1QMW | Yes |
| 1RPB | Yes |
| 1S6W | No |
| 1TER | No |
| 1UL2 | Yes |
| 1V6R | Yes |
| 1XGA | Yes |
| 1XGB | No |
